# Indication of family-specific DNA methylation patterns in developing oysters

**DOI:** 10.1101/012831

**Authors:** Claire E. Olson, Steven B. Roberts

## Abstract

**Background:** DNA methylation is an epigenetic modification that is ubiquitous across many eukaryotes, with variable patterns and functions across taxa. The roles of DNA methylation during invertebrate development remain enigmatic, especially regarding the inheritance and ontogenetic dynamics of methylation. In order to better understand to what degree DNA methylation patterns are heritable, variable between individuals, and changing during *Crassostrea gigas* development, we characterized the genome-wide methylome of *Crassostrea gigas* sperm and larvae from two full-sib families nested within a maternal half-sib family across developmental stages.

**Results:** Bisulfite treated DNA sequencing of *Crassostrea gigas* sperm and larvae at 72 hours post fertilization and 120 hours post fertilization revealed DNA methylation ranges from 15-18%. Our data suggest that DNA methylation patterns are inherited, as methylation patterns were more similar between the two sires and their offspring compared to methylations pattern differences among developmental stages. Loci differing between the two paternal full-sib families (189) were found throughout the genome but were concentrated in transposable elements. The proportion of differentially methylated loci among developmental stages was not significantly greater in any genomic region.

**Conclusions:** This study provides the first single-base pair resolution DNA methylomes for both oyster sperm and larval samples from multiple crosses. Assuming DNA methylation is introduced randomly, the predominance of differentially methylated loci between families within transposable elements could be associated with selection against altering methylation in gene bodies. For instance, differentially methylated loci in gene bodies could be lethal or deleterious, as they would alter gene expression. Another possibility is that differentially methylated loci may provide advantageous phenotypic variation by increasing transposable element mobility. Future research should focus on the relationship between epigenetic and genetic variation, and explore the possible relationship of DNA methylation and transposable element activity.

## Introduction

DNA methylation is an epigenetic modification that is ubiquitous across many eukaryotes, with variable patterns and functions across taxa. This epigenetic mechanism involves the addition of a methyl group to cytosines, usually in CpG dinucleotides, catalyzed by DNA methyltransferases. Epigenetic modifications such as DNA methylation can alter gene expression without modifying the underlying nucleotide sequence, and functions in mammals to suppress transcription through increased methylation in promoter regions (Bell and Felsenfeld, 2000). In mammals, DNA methylation is essential for development and differentiation of organs and tissues (Okano et al. 1999). Likewise, mutations of DNA methyltransferase in mammals may result in developmental delays and mortality (Li et al. 1992).

In contrast to the densely methylated mammalian genomes, several invertebrate species display low to intermediate levels of methylation. In invertebrates, it has been proposed that DNA methylation of genes may be associated with alternative splicing events (i.e. honey bee (Lyko et al. 2010) and *Nasonia* (Park et al. 2011)). Methylation of gene bodies has also been shown to have a positive relationship with transcriptional activity in oysters (Gavery and Roberts, 2013; Olson and Roberts, 2014a). Currently there is an incomplete understanding of the regulation of gene expression by DNA methylation in invertebrates, though it appears to be distinct from mechanisms observed in mammals and likely varies across species.

From the limited studies that have focused on invertebrates, research has shown that similar to mammals, DNA methylation has important regulatory functions during early development. For example, research on the honey bee *Apis mellifera* found DNA methylation to be abundant in the genome, with methylation being associated with altered gene expression resulting in bee caste differentiation (Elango et al. 2009; Kucharski et al. 2008). Furthermore, DNA methylation has been shown to regulate gene expression during *Octopus vulgaris* development, particularly during the first paralarval stage that includes significant morphological changes (Diaz-Freije et al. 2014). The first indication that methylation was an important regulator of development in *C. gigas* was by Riviere et al. (2013), who found treatment with 5-Aza-cytidine leads to developmental alterations and abnormal phenotypes in oysters.

Despite the essential role of methylation in development, there is limited information on individual variation in DNA methylation patterns among invertebrates and particularly how any methylation information might be passed on to offspring. Furthermore, we do not have a full understanding of ontogenetic changes in DNA methylation. In order to better understand to what degree DNA methylation patterns are heritable, variable between individuals, and changing during *C. gigas* development, we analyzed genome-wide DNA methylation in gametes and larval oysters (72 and 120 hours post-fertilization) from two full-sib families.

## Methods

### Experimental Design

Oysters (two males and a single female) were collected from Oakland Bay, South Puget Sound, WA. No ethical approval or permits were required to work with this species. Oysters were strip spawned by scoring the gonad tissue with a sterile razor blade and rinsing out gametes with sterile seawater. Oocytes were incubated for 30 minutes in sterile seawater and 2 million oocytes each were placed into two separate plastic containers. Sperm diluted in sterile seawater (1L) from each male were used to fertilize oocytes. Fertilization was confirmed by examining polar bodies in cells under a compound microscope.

Larvae were kept in static tanks (100L) up to 120 hours post-fertilization (hpf). Counts of oyster larvae were performed at 120 hpf to confirm normal development. Two samples for DNA methylation analyses were taken from sperm prior to fertilization, and larvae samples collected at days 72 hpf and 120 hpf. Larvae samples were taken by filtering on a 20μm screen. All samples were preserved in 95% ethanol. For simplicity the sperm and corresponding larvae samples are referred to as family #1 and family #3 based on paternity.

### Bisulfite treated DNA Sequencing (BS-Seq)

Genomic DNA was extracted using DNAzol according to the manufacturer’s protocol (Molecular Research Center, Inc., Cincinnati, OH). High molecular weight genomic DNA (6 ug per sample), was used to prepare six libraries for whole-genome bisulfite sequencing. Briefly, DNA was fragmented to an average length of 250 bp in an Adaptive Focused Acoustics (AFA) microtube using a Covaris S2 (Covaris Inc Woburn, MA) with the following settings: duty cycle 20%, intensity of 4.0, cycles per burst 200, for 60 seconds. Fragment size resulting from DNA shearing was confirmed by gel electrophoresis. Libraries were constructed using the Paired-End DNA Sample Prep Kit (Illumina, San Diego, CA) with standard protocols. Unmethylated Lambda DNA (0.5%) (Promega Co. Madison, WI) was added to the each sample prior to fragmentation and library construction to serve as a measure of bisulfite conversion efficiency. DNA was treated with sodium bisulfite using the EpiTect Bisulfite Kit (Qiagen, Valencia, CA) and 72 bp paired-end sequencing was performed on the Illumina HiSeq 2000 system. Library construction and sequencing was performed by the High Throughput Genomics Center (htSEQ, Seattle, WA).

Bisulfite sequencing reads from the six libraries were quality filtered to remove adapter sequences and separately mapped to the *Crassostrea gigas* genome (version GCA_000297895.1; Zhang et al. 2012) using Bisulfite Sequencing Mapping Program BSMAP v2.74 (Xi and Li 2009). Resulting alignment from mapping bisulfite treated reads was analyzed with *methratio*, a Python script that accompanies BSMAP. Parameters for *methratio* included reporting loci with zero methylation ratios (-z), combining CpG methylation ratios on both strands (-g) and only using unique mappings (-u). Only CpG loci covered by at least 3 sequenced reads were considered for further analysis. Data can be accessed and computational analysis performed as described via a GitHub repository (Olson and Roberts, 2014b). The IPython notebook in this repository includes all steps necessary for downloading and reproducing the analyses described in this manuscript.

### Global DNA Methylation Comparison

Whole-genome DNA methylation correlation and clustering were performed using the program methylKit 0.9.2 (Akalin et al. 2012) in R v3.0.3. Pairwise Pearson’s correlation coefficients scores were calculated based on the percent methylation profiles between all pairs of samples. Hierarchical clustering was performed using 1-Pearson’s correlation distance of the six methylation profiles.

### Differentially Methylated Loci

Differential methylation between the two full-sib families at each locus was determined using Fisher’s exact test in methylKit. A CpG locus was determined to be different between families when the difference in methylation ratio between families was more than 25% and *p*-value <0.01.

Ontogenetic changes in DNA methylation patterns were tested by three pairwise comparisons (Fisher’s exact tests in methylKit) between all developmental stages (sperm and 72 hpf larvae, sperm and 120 hpf larvae, and 72 hpf and 120 hpf larvae). Differentially methylated loci were identified as any CpG with greater than 25% and *p*-value <0.01 for any comparison.

All loci with significantly different methylation ratios across families and ontogenetic stages were characterized with respect to genomic features (intron, exon, promoter region, transposable element) using Bedtools (i.e., intersectBed) (Quinlan and Hall, 2010). Genomic features were developed and reported elsewhere (Gavery and Roberts 2013). Putative promoters were defined as 1kb regions upstream from transcription start sites. Transposable elements were identified using RepeatMasker, a program that screens and annotates interspersed repeats (Smit et al., 1996-2010). Specifically, RepeatProteinMask, was used with Repbase (Jurka et al. 2005), which contained 7,445 sequences. Sequence similarities from RepeatProteinMask were examined using WU-Blast (Lopez et al., 2003) to identify transposable elements which were included in the genome feature track. A total of 58,468 transposable elements identified based on sequence similarity. This genome feature track along with intron, exon, and promoter region feature track are all available via the IPython notebook that provides code and data used in this manuscript (Olson and Roberts, 2014b). A Chisquared test was performed to determine if the distribution of differentially methylated loci among genomic regions (intron, exon, promoter region, transposable elements) was significantly different to the distribution of all CpGs in the oyster genome.

## Results

### Bisulfite treated DNA Sequencing (BS-Seq)

Bisulfite treated DNA sequence reads are available (NCBI Sequence Read Archive: Study Accession Number SRP028178 - Accession Numbers SRX795174-SRX795179). Sodium bisulfite conversion efficiency was estimated to be 99.9% based on analysis of lambda phage DNA. The number of sequenced cytosines ranged from 2.6x10^7^ to 5.3x10^7^ across libraries. Using a 3x coverage threshold, most cytosines (75-78%) were determined to be unmethylated (methylation ratio = 0), while 15-18% of the CpG dinucleotides were methylated (methylation ratio 0.5) (data not shown).

### Genome-wide DNA Methylation Comparison

Relationships on a genome-wide scale were assessed by sample correlation and clustering. A total of 40,654 common loci were shared among all six samples and thus analyzed. Methylation ratios were highly correlated between sperm and respective progeny, with a pair-wise Pearson’s correlation coefficient (*r*) of 0.84 for both families. These similarities within families are also evident in hierarchical clustering (Figure 1), where both male gamete samples were more similar in their methylation profiles to their respective 120 hpf larvae.

**Figure 1.**
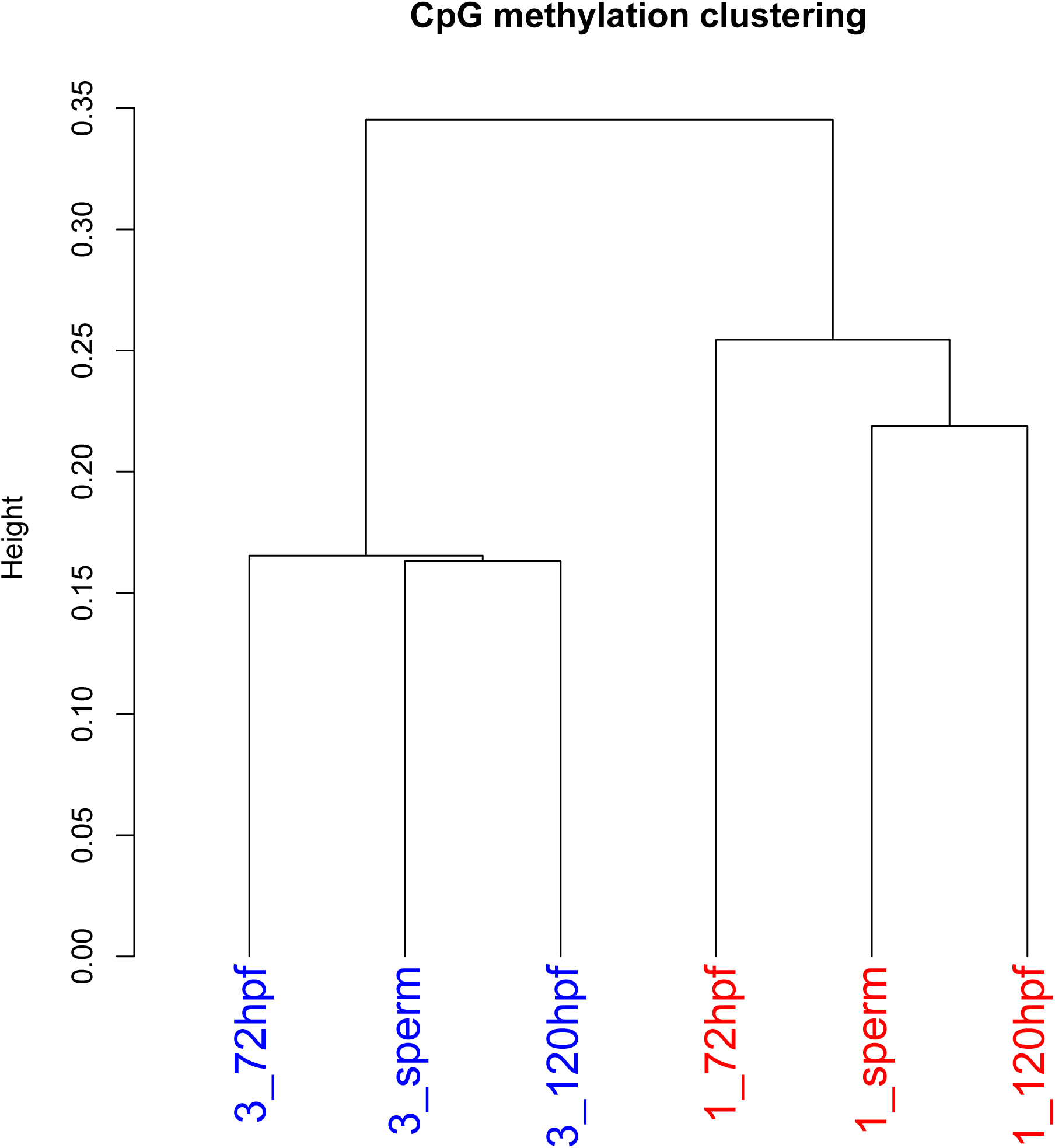
Dendogram of the male sperm and oyster larvae genome-wide methylation profiles using Pearson’s correlation distance. Numeric prefix refers to family.

### Family-specific and Developmental Differences

A total of 189 differentially methylated loci (DMLs) were identified between the two full-sib families. Of these, 99 were found to overlap with a defined genomic region (exon, intron, promoter region, transposable element) (Figure 2). Most CpG loci with different methylation ratios among oyster families were in introns. However, compared to the distribution of CpG dinucleotides in the oyster genome, the proportion of differentially methylated loci within transposable elements was significantly higher than expected (*χ*^2^= 22.17, df= 1, p < 0.0001).

**Figure 2.**
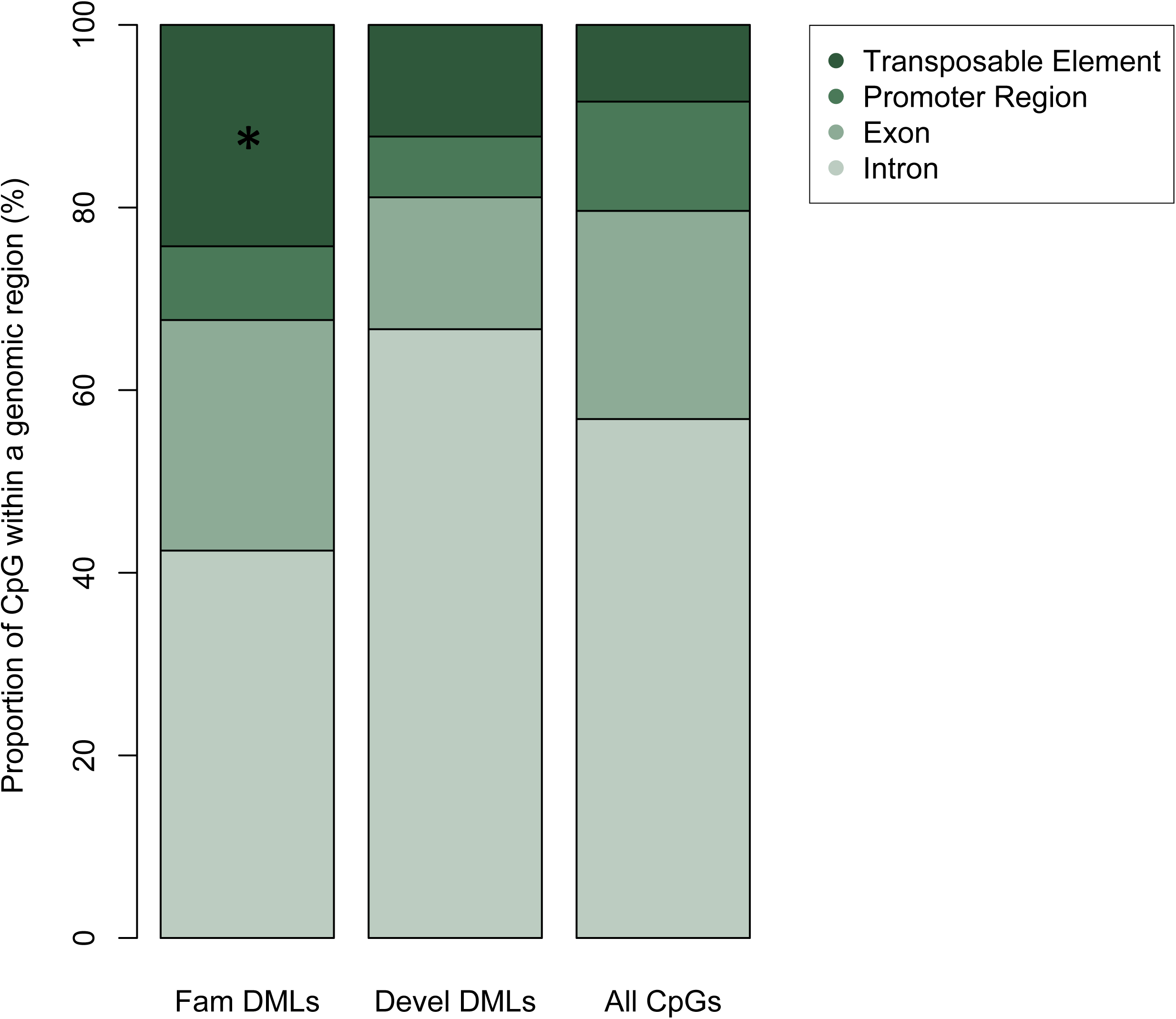
Proportion of family-specific differentially methylated loci, developmentally different differentially methylated loci, and all CpGs in the oyster genome based on genomic region. The proportion of loci located within a genomic region (Intron, Exon, Promoter Region, Transposable Element) for differentially methylated loci between families (n= 99), differentially methylated loci among the developmental stages (n= 90), and all CpGs in the oyster genome (n= 10035701) are displayed. An asterisk indicates a statistically different distribution relative to the distribution of all CpGs in the oyster genome.

A total of 160 CpG loci showed differences in methylation ratios among developmental stages. Of these loci, 90 were within defined genomic regions (Figure 2). The proportion of differentially methylated loci was not significantly greater in any genomic region analyzed.

## Discussion

This study provides the first single-base pair resolution DNA methylomes for both oyster sperm and larval samples from multiple crosses. This research not only provides new information on DNA methylation patterns during oyster development, but also examines its inheritance and changes during early development. Interestingly, our research indicates that epigenetic patterns may differ among oyster families.

Methylation levels in oyster sperm and larvae samples ranged from 15-18% with interspersed regions of both methylated and unmethylated DNA in both male gamete and larval samples. This proportion of CpG methylation falls within the range of that previously reported for oyster male gonad tissue (Olson and Roberts, 2014a) and oyster gill tissue (Gavery and Roberts, 2013). Overall methylation levels are also comparable to those reported among multiple developmental stages of the Pearl oyster *Pinctada fucata* (Li et al. 2014). These findings indicate that overall genome methylation in *C. gigas* is at an intermediate level and suggests that DNA methylation levels do not significantly vary between multiple cell types and life history stages. This is similar to what has been described in global 5-methylcytosine content during different stages of *Ciona intestinalis* development (Suzuki et al. 2013). However, it contrasts with research on mammals and vertebrates, which exhibit the presence of tissue and developmental stage specific methylation profiles, as that seen in zebrafish (McGaughey et al. 2014).

Genome-wide comparisons indicated higher similarity of methylation patterns between oyster families than between developmental stages, suggesting that DNA methylation patterns are inherited. If epigenetic marks are indeed heritable, this mechanism has significant implications for selection. It has been proposed that epigenetic variation may compensate for a decrease in genetic variation in species such as sparrows (Shrey et al. 2012). While outside the scope of the current study, an assessment of relationships between genetic and epigenetic variation is critical. Several studies have examined epigenetic differentiation in vertebrate and plant populations experiencing different environments, indicating evidence for divergent selection in these species (Liu et al. 2012; Herrera and Bazaga 2010). Few studies have focused on invertebrates, though Jiang et al. (2013a) investigated the relationship between genetic and epigenetic variations in two groups of *C. gigas*, a base stock domesticated population and third generation mass selection population. This study demonstrated genetic differentiation between the base population and third generation mass selection populations of oysters, but did not find overall epigenetic variation (Jiang et al. 2013a). Nevertheless, a significant correlation was observed between genetic and epigenetic profiles, with few individuals having similar genetic but distinct epigenetic profiles (Jiang et al. 2013a). Regardless, if epigenetic variation is independent of genotype, mechanisms involved in epigenetic inheritance are not fully understood.

Differences in genome-wide methylation ratios between full-sib families nested within a maternal half sib family suggested paternal inheritance of DNA methylation patterns. These results are similar to what has been seen in zebrafish where embryos inherit the methylation profile of sperm rather than oocyte (Jiang et al. 2013b, Potok et al. 2013). In contrast, methylation ratios in Pearl oysters are mainly influenced by oocytes, rather than sperm (Li et al. 2014), possibly due to maternal influences on DNA methylation patterns in early larvae, while later stages attain methylation patterns similar to sperm.

Differentially methylated loci across families were distributed throughout the genome, though a higher proportion was found in transposable elements. This concentration of methylation in transposable elements may be due to selection against altering methylation in functionally important parts of the genome. For instance, many differentially methylated loci in gene bodies could be lethal or deleterious as they would alter gene expression. It should be noted that the role of DNA methylation in regulating genome activity in *C. gigas* is still unclear. However, it has been suggested that elevated methylation decreases spurious transcription of housekeeping genes and limited methylation in inducible genes facilitates multiple transcriptional opportunities (Roberts and Gavery, 2012). In other words, DNA methylation patterns in gene bodies may have evolved over time based on gene function to fit the needs of organisms in highly variable environments, and random changes in these patterns could be detrimental. Furthermore, we suggest that random variations in methylation within transposable elements may have a relatively higher chance of persisting than elsewhere in the genome. Transposable elements are mobile DNA sequences that may be methylated in many species to silence activity (Yoder et al. 1997; Liu and Schmid 1993). Limited information is available about the methylation status of transposable elements in other invertebrate species, but the available studies suggest that transposons are generally unmethylated and contain similar levels of methylation to neighboring DNA (Suzuki and Bird, 2008). This is in agreement with our previous research, which showed limited DNA methylation in transposable elements in oyster male gamete tissue (Olson and Roberts, 2014a). Assuming that transposable element activity is less critical to survival than coding gene activity, differentially methylated loci in transposable elements may be less likely to have negative selective effects. On the other hand, differentially methylated loci may also provide advantageous phenotypic variation by increasing transposable element mobility. However, such selection hypotheses assume that methylation is introduced randomly, something we do not have evidence for.

Interestingly, we did not observe a high proportion of differentially methylated loci among promoter regions, as would be expected if promoter methylation was regulating gene expression to play a role in oyster development. Recent research has found that DNA methylation of promoter regions specifically reduces expression of *Hox* genes during oyster development (Riviere et al. 2013). Considerable stage-specific differences in total methylation levels during oyster early development indicated that DNA methylation plays a crucial role in oyster embryogenesis (Riviere et al. 2013). We previously found variation in expression levels depending on the level of promoter region methylation (Olson and Roberts, 2014a). Surprisingly, we did not observe any dramatic differences in overall methylation levels during oyster development, nor higher methylation of promoter regions. This discrepancy is likely due to the analysis of different ontogenetic stages, as Riviere et al. (2013) examined the first 24 hours post-fertilization, and our first larval sample was taken at 72 hpf. It is also possible that only a subset of genes are transcriptionally controlled via DNA methylation and our global approach masked the ability to see differences.

## Conclusions

This research suggests epigenetic inheritance as DNA methylation patterns were similar between males and their offspring and differed between oyster families. Interestingly, we found a high proportion of family-specific methylation patterns within transposable elements. Future research should focus on the relationship between epigenetic and genetic variation, and explore the possible relationship of DNA methylation and transposable element activity.

## Acknowledgements

This work was supported by a National Science Foundation award (Grant Number 1158119) to SR.

## Abbreviations/Definitions

BAFA: Adaptive Focused Acoustics
BS-Seq: Bisulfite Sequencing
DMLs: Differentially Methylated Loci
hpf: hours post-fertilization

## Competing Interests

The authors declare that they have no competing interests.

## Author Contributions

CO carried out the experimental trial, molecular work, and bioinformatic analyses. CO and SR participated in the design of the study and drafting of the manuscript. All authors read and approved the final manuscript.

